# Whole genome sequencing analysis of the cardiometabolic proteome

**DOI:** 10.1101/854752

**Authors:** Arthur Gilly, Young-Chan Park, Grace Png, Andrei Barysenka, Iris Fischer, Thea Bjornland, Lorraine Southam, Daniel Suveges, Sonja Neumeyer, N. William Rayner, Emmanouil Tsafantakis, Maria Karaleftheri, George Dedoussis, Eleftheria Zeggini

**Affiliations:** Institute of Translational Genomics, Helmholtz Zentrum München – German Research Center for Environmental Health, Neuherberg, Germany; Wellcome Sanger Institute, Wellcome Genome Campus, Hinxton CB10 1SA, UK; University of Cambridge, Cambridge, UK; Department of Mathematical Sciences, Norwegian University of Science and Technology, NO-7491 Trondheim, Norway; Wellcome Centre for Human Genetics, Oxford, UK; European Bioinformatics Institute, Wellcome Genome Campus, Hinxton CB10 1SH, UK; Wellcome Centre for Human Genetics, Nuffield Department of Medicine, University of Oxford, Oxford, UK; Oxford Centre for Diabetes, Endocrinology and Metabolism, Radcliffe Department of Medicine, University of Oxford, Oxford, UK; Anogia Medical Centre, Anogia, Greece; Echinos Medical Centre, Echinos, Greece; Department of Nutrition and Dietetics, School of Health Science and Education, Harokopio University of Athens, Greece

## Abstract

The human proteome is a crucial intermediate between complex diseases and their genetic and environmental components, and an important source of drug development targets and biomarkers. Here, we comprehensively assess the genetic architecture of 257 circulating protein biomarkers of cardiometabolic relevance through high-depth (22.5x) whole-genome sequencing (WGS) in 1,328 individuals. We discover 131 independent sequence variant associations (*P*<7.45×10^−11^) across the allele frequency spectrum, all of which replicate in an independent cohort (n=1,605, 18.4x WGS). We identify for the first time replicating evidence for rare-variant *cis*-acting protein quantitative trait loci for five genes, involving both coding and non-coding variation. We construct and validate polygenic scores that explain up to 45% of protein level variation. We find causal links between protein levels and disease risk, identifying high-value biomarkers and drug development targets.

---

Cardiometabolic diseases are a leading cause of death and continue to rise in prevalence across global populations. Biomarkers can help improve early detection and diagnosis, and can lead to better clinical outcomes through improved and timely interventions. To comprehensively characterise the genetic underpinning of protein biomarker levels, disentangle correlation from causation, and identify opportunities for new therapeutic target discovery and predictive modeling, we first assessed the association between 257 cardiometabolic disease-related serum protein levels^1^ and 13,419,876 single nucleotide variants (SNVs) in a population-based cohort (MANOLIS) with deep whole genome sequence data.

We identify 116 protein quantitative trait loci (pQTLs) reaching study-wide significance (*P*<7.45×10^−11^) (Supplementary Table 1, Figure 1). Thirty-two (27%) of these are driven by multiple independent variants (between two and seven per locus, giving rise to a total of 164 independently-associated variants) illustrating complex allelic architecture at pQTLs (Supplementary Figure 1). We find replicating evidence for association (*P*<0.000305) across 131 out of 159 variants (82%) present in an independent, whole genome-sequenced population-based cohort with the same serum biomarker measurements (Pomak^2^) (n=1,605, 18.4x WGS, Supplementary Figure 2). We find that these robustly-replicating loci explain up to 47.7% of protein level variance, and on average more (one-sided Mann-Whitney-U test, *P=*3.42×10^−13^) than for 37 other, non-proteomic quantitative traits measured in the same individuals (Supplementary Figure 3). This exemplifies how the study of blood biomarkers can powerfully capture the heritable component of biological processes underpinning such disease-relevant quantitative traits. Eighty of the associated variants display significant allele frequency differences between the isolated population studied here and large reference populations (Supplementary Table 2), 53 of which have increased frequencies in MANOLIS (66%, *P*=0.002, one-sided 1-sample proportion test). In particular, 14 associated variants display a frequency increase of more than 5-fold, highlighting the advantage of using isolated populations in associations of protein levels.

**Figure 1:**
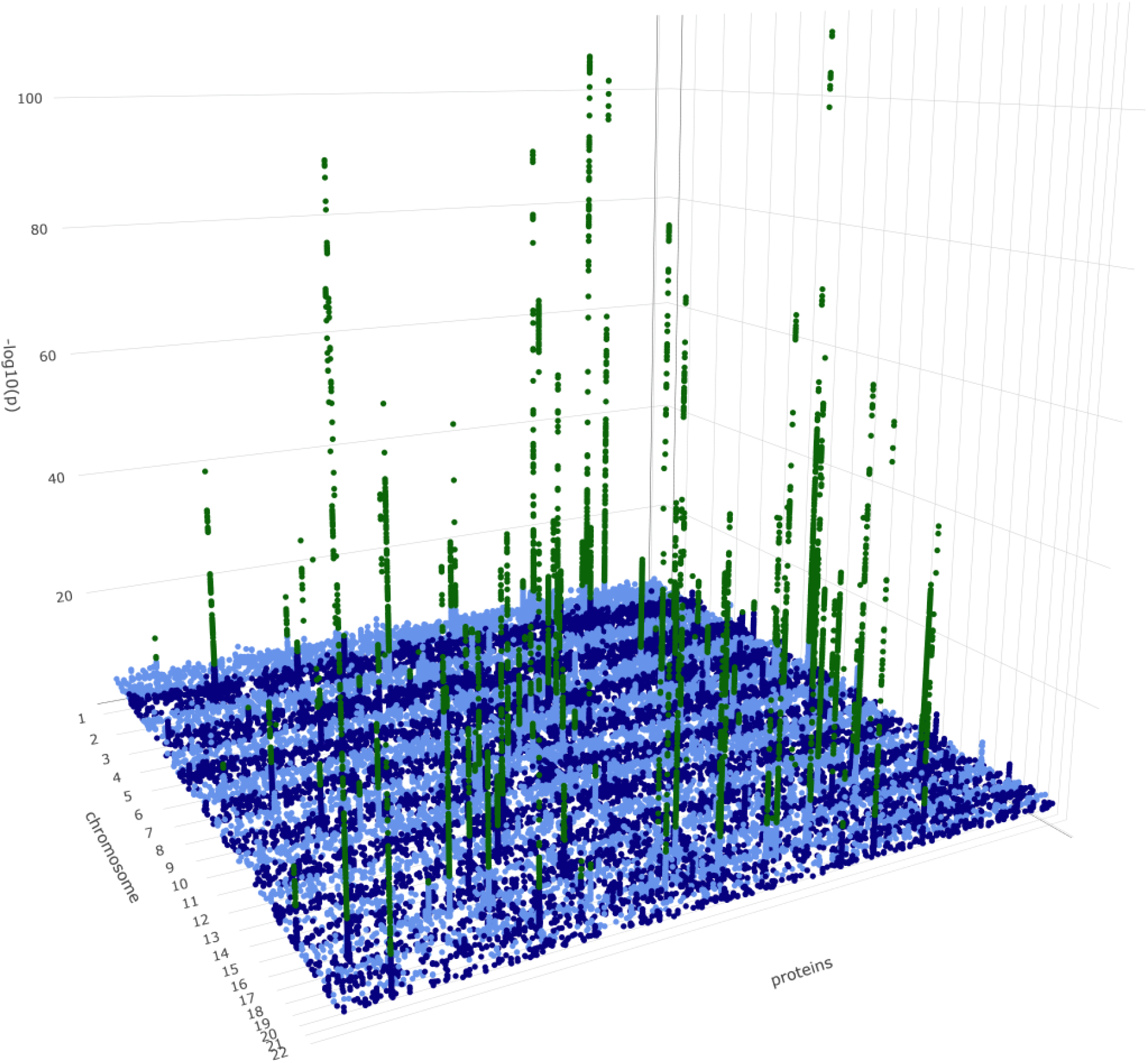
Genome-wide association signals across all tested proteins. For clarity, variants with *P*>1×10^−5^ are not represented in the figure. Variants with *P*<7.45×10^−11^ are plotted in green.

117 (90%) of these 131 reproducibly-associated variants are common (minor allele frequency (MAF) >5%), 13 are low-frequency (MAF 1-5%) and 1 is rare (MAF<1%) (Figure 2). One hundred of the associated variants (76%) are located within 1Mb of the gene encoding the respective protein (i.e. in *cis-*pQTLs), and 31 (24%) are in *trans*-pQTLs (Figure 2). 57 *cis*-associated variants are located within the boundaries of the respective gene, and the remaining 43 are at a median distance of 13.8kb (max. 920 kb) (Supplementary Figure 4).

**Figure 2:**
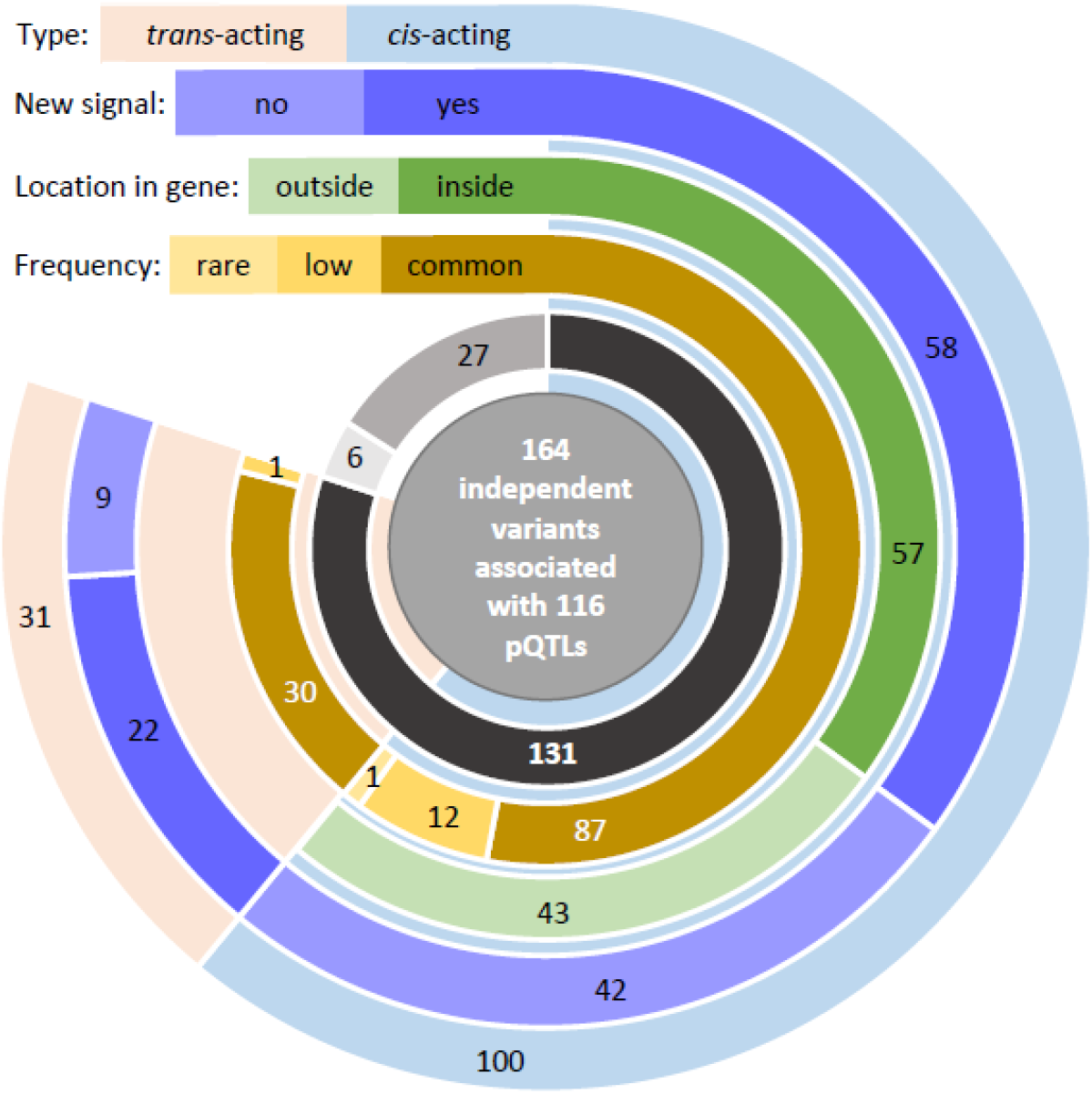
Characteristics of independently contributing pQTL variants. The innermost circle represents replication status: dark grey for variants that replicate, medium grey for variants that do not replicate and light grey for variants for which no proxy was found in the Pomak dataset.

Thirty-two out of 72 *cis*-pQTLs (44%) discovered in this cohort have either not previously been reported in protein-level GWAS (novel loci), or harbour variants conditionally independent of all previously-reported associations (novel variants at known loci) (Supplementary Table 1).

We identify 38 variants in 35 *trans* loci associated with 32 proteins; 18 of these variants both have not been previously reported (Supplementary Text) in protein-level GWAS and replicate in the Pomak population. We find the overall replication rate to be similar for *trans*-(81%) and *cis*-associated (79%) variants. Of the replicating 31 *trans*-acting variants, 30 are common and one is low-frequency. We identify *trans*-pQTL signals for seven receptor/ligand pairs with experimental evidence of physical interaction and well-established synergistic roles in downstream pathways^3–9^.

To enhance our understanding of rare variant (RV) contribution to serum protein biomarker levels, we performed gene-based burden analysis across coding and non-coding sequence variation (Methods). We identify for the first time 6 study-wide significant (*P*<7.45×10^−11^) *cis*-RV-pQTLs (Figure 3, Supplementary Figure 5), in the *ACP6* (lysophosphatidic acid phosphatase type 6, *P*_meta-analysis_=3.17×10^−97^), *PON3* (paraoxonase 3, *P*_meta-analysis_=7.42×10^−86^), *IL1RL1* (interleukin 1 receptor like 1, *P*_meta-analysis_=2.15×10^−58^), *DPP7* (dipeptidyl peptidase 7, *P*_meta-analysis_=2.71×10^−36^), *CTSO* (cathepsin O, *P*_meta-analysis_=2.27×10^−33^) and *GRN* (progranulin, *P*_MANOLIS_=3.16×10^−12^) genes. All except the *GRN* burden signal replicate in the Pomak cohort. The *GRN* RV-pQTL is driven by the novel splice donor variant chr17:44349552 (minor allele count MAC=4) and the 5’UTR variant rs563336550 (MAF=1.7%) in MANOLIS, the latter showing a 17-fold increase in frequency in MANOLIS compared to gnomAD non-Finnish Europeans (MAF=0.1%), and more than 2000-fold compared to TOPMed (MAF=0.00079%).

**Figure 3:**
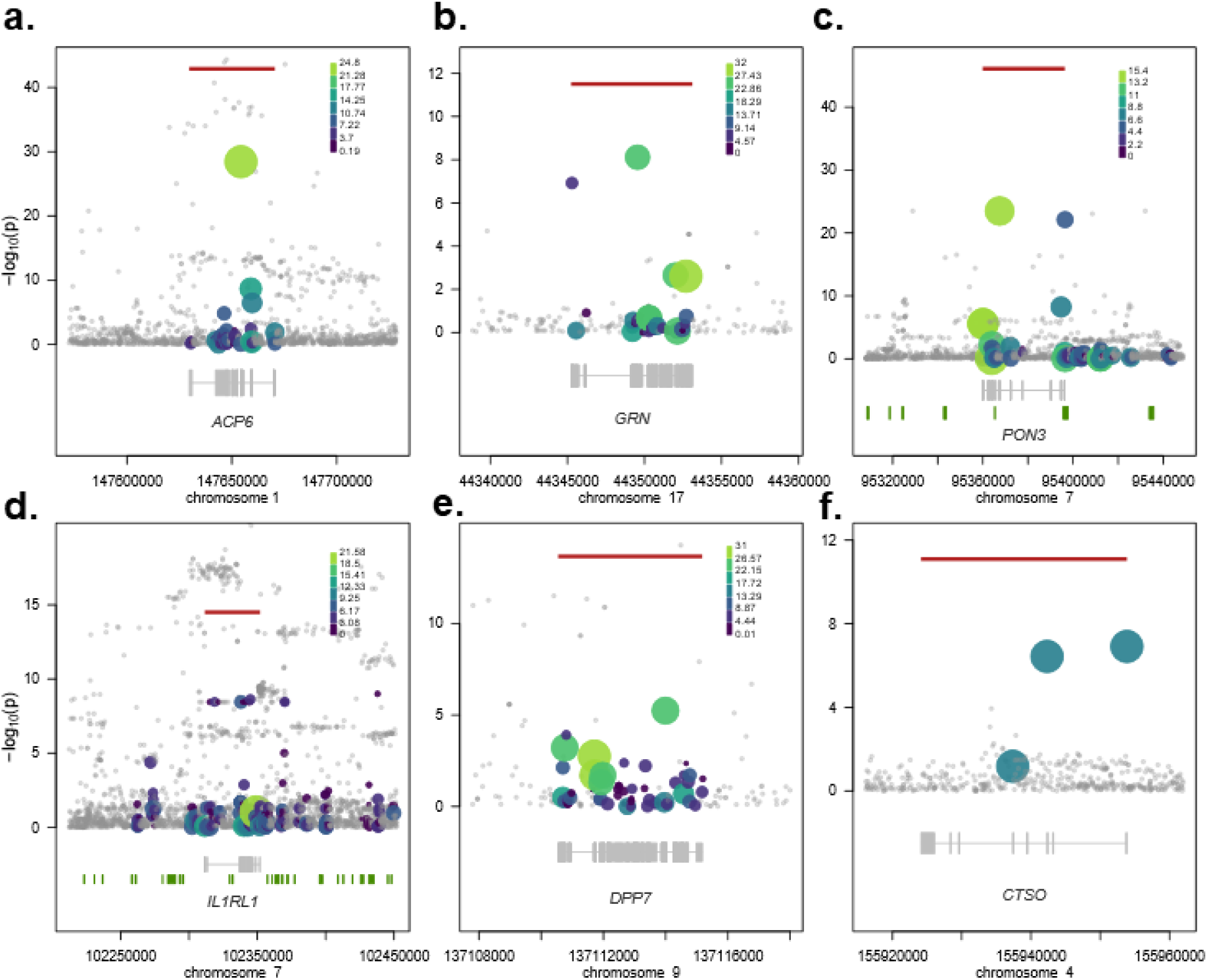
Rare variant pQTLs. Rare variant burden signals detected in this study -the most significant burden per gene is displayed. Circles denote the sequence variants identified in the region. Genes are denoted in gray below the regional association plots; bars represent exons across all transcripts. Horizontal red lines indicate the −log10 of the burden signal p-value, with size and colour of circles proportional to the weighting scheme used (CADD or Eigen). Grey circles denote variants not included in the burden. Details on variants included in each burden are given in Supplementary Table 14.

We find that rare regulatory variants are major contributors to some of these burdens. For example, one of the two variants driving the PON3 *cis*-RV-pQTL resides in promoter ENSR00000215353 and transcription factor binding site ENSR00000832511, and is associated with a decrease in PON3 levels (rs149867961, MAF=3.1%, effect size β=−1.18 in units of standard deviation, standard error σ=0.113, *P*=7.58×10^−23^). The other contributing variant, rs772677677 (MAF=1.9%, β=−1.55, σ=0.143, *P*=3.15×10^−24^), is a missense variant with a substantially increased frequency in MANOLIS (MAF=1.9% compared to 0.00264% in gnomAD), also associated with a decrease in PON3 levels. PON3 (paraoxonase 3) inhibits the oxidation of low-density lipoprotein (LDL), an effect that slows atherosclerosis progression^10^. These findings illustrate the contribution of rare variants to the heritability of proteomic traits, and that this contribution is partly mediated through *cis*-RV-pQTLs.

To detect proteins that may play a causal role in cardiometabolic disease onset or progression, we performed two-sample Mendelian randomization analysis across 93 proteins with study-wide significant signals here and 193 diseases and traits from UK Biobank and other large consortial datasets (Methods, Supplementary Table 3). We identify significant (FDR<0.05) associations involving 48 proteins and 75 phenotypes (Supplementary Table 4, Figure 4). For 13 of these proteins, pQTL SNPs had both a lowering effect on circulating levels and a protective effect against at least one disease (Supplementary Table 5), suggesting potential antibody-based approaches for therapeutic benefit.

**Figure 4:**
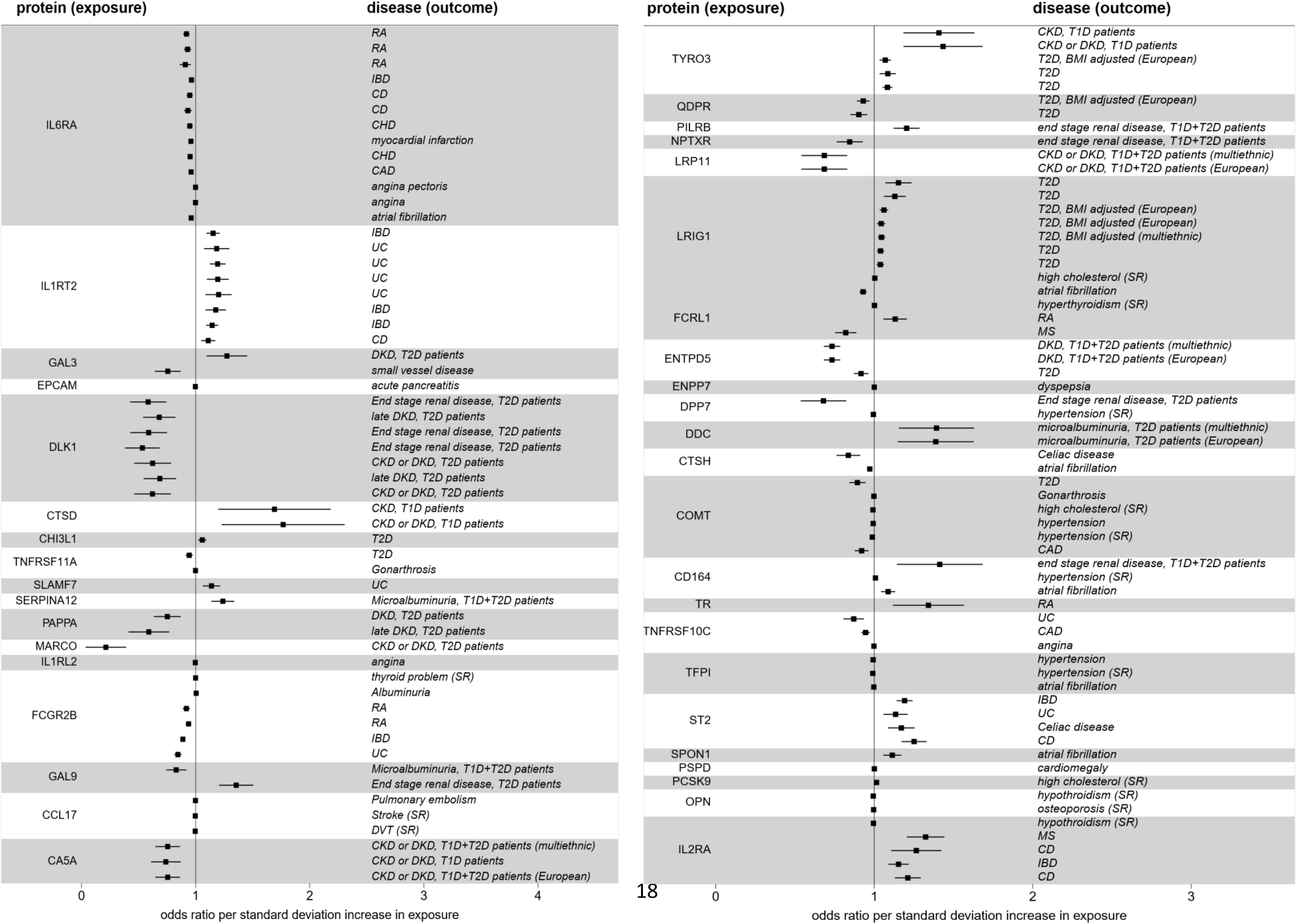
Significant causal protein-disease associations identified through two-sample Mendelian randomisation. Protein (exposure) names are indicated on the left, diseases (outcomes) on the right. Identical disease names for a given protein indicate a MR signal replicating across multiple studies of the same disease; further details and causal associations with quantitative traits are displayed in Supplementary Table 4. RA: rheumatoid arthritis, IBD: inflammatory bowel disease, CD: Crohn’s disease, CHD: Coronary heart disease, CAD: Coronary artery disease, UC: ulcerative colitis, DKD: diabetic kidney disease, T2D: type 2 diabetes, T1D: type 1 diabetes, CKD: chronic kidney disease, MS: multiple sclerosis.

Providing proof of principle, we find evidence of established links between dysregulated protein levels and common diseases, such as an inverse causal correlation between PCSK9 levels and hypercholesterolemia (*P*=1.00×10^−10^, *P*_*FDR*_=9.46×10^−8^), and between osteopontin levels and risk of osteoporosis (*P*=6.49×10^−5^, *P*_*FDR*_=0.011) and hypothyroidism (*P*=1.2×10^−4^, *P*_*FDR*_=0.016). Similarly, we find decreased levels of IL1RL1 and IL1RT2, both proteins involved in autoimmunity and inflammation-related disorders^11^, to be causally linked to risk of autoinflammatory bowel diseases (Supplementary Table 4).

We further identify new evidence for disease-mediating roles for proteins circulating in the periphery. For example, rs2306272, a missense *cis-*pQTL, is associated with decreased LRIG1 (leucine rich repeats and immunoglobulin like domains 1) levels (MAF=31%, meta-analysis β=−0.754, σ=0.0261, *P=*1.50×10^−183^), and is causally associated with reduced risk of atrial fibrillation (*P*=5.23×10^−11^, *P*_*FDR*_=5.32×10^−8^) and lower BMI (*P*=2.72×10^−9^, *P*_*FDR*_=1.89×10^−6^), and with increased risk of type 2 diabetes (*P*=4.70×10^−5^, *P*_*FDR*_=8.87×10^−3^) and self-reported hypercholesterolemia (*P*=4.61×10^−4^, P_FDR_=0.043) (Figure 4). LRIG1 is a transmembrane protein that acts as a feedback negative regulator of signaling by receptor tyrosine kinases. Variants in *LRIG1* have previously been associated with atrial fibrillation^12,13^, and the identified pQTL co-localises with previous pulse rate (posterior probability of colocalisation *P*_4_=0.939) and QRS duration (*P*_4_=0.998) loci. Mouse knockout models of *LRIG1* exhibit decreased body weight and fat^14^.

Notably, we find evidence for a genetic link between *PRG2* intronic variant rs10642232 and decreased levels of PAPPA (pregnancy-associated plasma protein-A) (β=−0.299, σ=0.0320, *P*=1.06×10^−20^). A previous association exists at rs140000161^15^, however this variant is not found in either of our cohorts. PAPPA is a metalloproteinase involved in normal and pathological insulin-like growth factor (IGF) physiology. *PRG2* codes for eosinophil granule major basic protein, which reduces PAPPA activity by interacting with it to form a complex^16^. PAPPA is a specific protease targeting IGFBP4 (IGF binding protein 4) in the presence of IGF. IGFBP4 inhibits IGF binding with its receptor, and PAPPA promotes IGF activity^17^. rs10642232 is a *PRG2*-decreasing eQTL in multiple tissues. We find reduced PAPPA levels to be causally associated with decreased risk of diabetic kidney disease in T2D patients (*P*=2.63×10^−4^, P_FDR_=0.0304). IGF activity is enhanced in early diabetic nephropathy, whereas IGF resistance is found in chronic kidney failure^18^. Animal knockouts of PAPPA exhibit decreased body weight and length, type 2 diabetes and hypercholesterolemia. Our results suggest that PAPPA and its inhibition by PRG2 within the IGF system may play a role in the pathogenesis and progression of diabetic kidney disease in T2D patients. These findings are consistent with the reported lower incidence of diabetic complications in this isolated Cretan population^19^.

Further, we find that *cis*-acting variants decreasing levels of ENTPD5 (rs73301485, MAF=7.5%, β=−0.637, σ=0.0487, *P*=4.59×10^−39^; rs140111715, MAF=3.7%, β=−0.778, σ=0.0563, *P*=2.29×10^−43^) are causally associated with lower risk of type 2 diabetes (*P*=3.56×10^−4^, *P*_*FDR*_=0.037) and diabetic kidney disease (*P*=2.61×10^−19^, *P*_*FDR*_=4.93×10^−16^). ENTPD5 (ectonucleoside triphosphate diphosphohydrolase 5) promotes glycolysis in proliferating cells in response to phosphoinositide 3-kinase (PI3K) signaling and is primarily expressed in the liver, kidney, intestine, prostate and bladder, and rs73301485 is an eQTL in multiple tissues. Mouse knockout models show decreased body weight, hypoglycemia, decreased cholesterol and triglycerides^20^. Small-molecule screens have recently identified several ENTPD5 inhibitors^21^ that warrant investigation for their effect on type 2 diabetes and diabetic complications.

We find that genome-wide polygenic scores calculated in MANOLIS can predict up to 45.5% of protein variance in the independent Pomak dataset, despite a low average predictive performance (median r^2^=0.026, Supplementary Figure 6). The polygenic score architecture observed within the power parameters of this study, indicates the involvement of a small number of strongly-associated common variants, and a smaller contribution for rare and low-frequency variants. Notably, both the discovery and test datasets stem from individuals of European ancestry; further studies in global populations will be required to assess the transferability of these polygenic scores.

Polygenic prediction of the cardiometabolic proteome can lead to the identification of potential biomarkers through correlation with disease states. We performed logistic regression of 47 proteins with polygenic scores that had achieved a predictive value of r^2^>0.05 in Pomak, on 86 indications in UK Biobank, adjusted for genetic principal components, clinical and lifestyle factors. We find that the scores for GRN (progranulin), CHI3L1 (chitinase 3 like 1) and PECAM1 (platelet and endothelial cell adhesion molecule 1) levels are significant predictors of disease status (Wald test *P*<1.66×10^−5^) across a range of cardiometabolic traits (Supplementary Table 6). The progranulin level polygenic score is correlated with increased risk of hypercholesterolemia, and is driven by a single association in *CELSR2*-*SORT1*, an established risk locus for lipid disorders^22^. In a joint predictive model for high cholesterol, inclusion of polygenic scores for CHI3L1 and PECAM1 levels results in a significant increase of the model accuracy compared to the clinical and lifestyle covariates-only model (Supplementary Text, Likelihood Ratio Test *P*=7.24×10^−27^). We find that an elastic net model agnostically selects the same three polygenic scores in a full-proteome analysis of high-cholesterol, confirming their contribution and demonstrating the value of including proteomics scores in predictive models of disease risk.

In summary, using whole-genome sequencing, we identify robustly-replicating *cis*- and *trans*-pQTLs, and show for the first time that burdens of rare variants contribute to the genetic architecture of protein biomarker levels. We show that incorporating information on this genetic contribution leads to improvement in clinical risk models for cardiovascular disease. Identification of causal contributions of the cardiometabolic proteome to the risk of multiple chronic diseases can present opportunities for new therapeutic target discovery and predictive modeling to accelerate precision medicine.

## Methods

### Sequencing and variant calling

Genomic DNA (500 ng) from 1482 samples was subjected to standard Illumina paired-end DNA library construction. Libraries underwent DNA sequencing using the HiSeqX platform (Illumina) according to manufacturer’s instructions. Variant calling was performed after filtering out contamination, spatial artifacts and duplicates, according to the Genome Analysis Toolkit (GATK) v. 3.5-0-g36282e4 Best Practices. The sequencing and variant calling is described in detail in a previous publication^23^.

### Proteomics

The serum levels of 276 unique proteins in 1,407 MANOLIS samples from three Olink panels - CVDII, CVDIII and Metabolism - were measured using Olink’s proximity extension assay (PEA) technology^1^. Briefly, for each assay, the binding of a unique pair of oligonucleotide-labelled antibody probes to the protein of interest results in the hybridisation of the complementary oligonucleotides, which triggers extension of by DNA polymerase. DNA barcodes unique to each protein are then amplified and quantified using microfluidic real-time qPCR. Measurements were given in a natural logarithmic scale in Normalised Protein eXpression (NPX) levels, a relative quantification unit. NPX is derived by first adjusting the qPCR Ct values by an extension control, followed by an inter-plate control and a correction factor predetermined by a negative control signal. This is followed by intensity normalisation, where values for each assay are centered around its median across plates to adjust for inter-plate technical variation. Further details on the internal and external controls used can be found at http://www.olink.com. Additionally, a lower limit of detection (LOD) value is determined for each protein based on the negative control signal plus three standard deviations. For our samples, NPX values that fall below the LOD were set to missing.

We adjusted all phenotypes using a linear regression for age, age squared, sex, plate number, and per-sample mean NPX value across all assays, followed by normalisation of the residuals. We also adjusted for season, given the observed annual variability of some circulating protein levels. Given the dry Mediterranean climate of Crete, we define season of collection as hot summer or mild winter. Plate effects are partially offset by the median-centering implemented by Olink. MANOLIS samples were plated in the order of sample collection, which results in plate and season information to be largely correlated.

### Quality control

We excluded 18 proteins across all panels with missingness or below-LOD proportion greater than 40%. BNP was measured across all three panels, and was excluded due to high missingness in CVDII. In total, we therefore excluded 19 protein measurements (total analysed 257, Supplementary Table 7). 26, 2 and 14 samples failed vendor QC and were excluded from CVDII, III and META, respectively. 42 samples were excluded due to missing age.

Sequencing data quality control has been described before^23^. Briefly, Variant-level QC was performed using the Variant Quality Score Recalibration tool (VQSR) from the Genome Analysis Toolkit (GATK) v. 3.5-0-g36282e4. Sample-level QC was performed by comparing genotypes with chip data in the same samples. Four individuals failed sex checks, 8 samples had low concordance with chip data, 11 samples were duplicates, and 12 samples displayed traces of contamination. As contamination and sex mismatches were correlated, a total of 25 individuals were excluded (*n* = 1457). Variants were further filtered using the Hardy-Weinberg equilibrium test at *P* = 1.0 × 10^−5^. We filtered out 14% of variants with call rates < 99%.

### Single-point association

We carry out single-point association using the linear mixed model implemented in GEMMA ^24^. We use an empirical relatedness matrix calculated on a LD-pruned set of low-frequency and common variants (MAF>1%) that pass the Hardy-Weinberg equilibrium test (*P*<1×10^−5^). We further filter out variants with missingness higher than 1% and MAC <10. 5 proteins were excluded due to having a genomic control λ_GC_<0.97 orλ_GC_>1.05 after association (Supplementary Table 7). 123 signals were extracted using the PeakPlotter software (https://github.com/hmgu-itg/peakplotter), which is based on a combination of distance-based and LD-based pruning. We extracted independent SNV at each associated locus using an approximate conditional and joint stepwise model selection analysis as implemented in GCTA-COJO^25^. To avoid overfitting when too many predictors are included in the model, we perform LD-based clumping using Plink v.1.9, based on an r^2^ value of 0.1 and a window of 1Mb prior to the GCTA-COJO analysis^26^. The extended linkage disequilibrium (LD) present within MANOLIS can cause very large peaks to be broken up into several signals. We identified and manually investigated 5 regions where multiple peaks were present, reducing the number of independent signals to 116 and the number of conditionally independent variants to 164. *Cis-*acting protein-altering variants may result in false-positive associations due to epitope effects. While exact quantification of such effects would have required comparison using proteomic measurements from an alternative assay method, we note that only 11 of these 100 replicating *cis-*acting variants have a potentially protein-truncating effect.

### Rare variant association

For rare variant association, we apply a MAF filter of 5% and a missingness filter of 1%. We use the linear mixed model extension of SKAT-O implemented in MONSTER^27^, using the MUMMY wrapper^23^ (https://github.com/hmgu-itg/burden_testing). Following our previously-reported analysis strategy^23^, we test for rare variant burden association on a gene-by-gene basis: firstly, restricting burdens to coding variants with Ensembl most severe consequence stronger than missense; secondly, including all coding variants weighted by CADD^28^; thirdly, including exon and regulatory variants using the phred-scaled Eigen score^29^; and, finally, regulatory variants only weighted by Eigen. Regulatory regions are linked to a gene if they overlap an Ensembl-documented eQTL for that gene in any tissue. A gene-pair was taken forward for quality control (QC) if it was significant in any of the four analyses. 17 signals passed this threshold. Because rare variant LD blocks can extend over long distances and capture overlapping common associations, we manually inspect LD blocks through the plotburden software (https://github.com/hmgu-itg/plotburden), and discard signals involving variants in LD with nearby *cis* ones. For each remaining signal, we then re-run the burden analysis conditional on the genotypes of the variant with the lowest single-point p-value that was previously included in the burden, so as to only consider signals arising from at least two distinct variants. 6 RV-pQTL signals pass this quality control procedure (Supplementary Figure 5).

### Replication

We performed replication in 1,605 samples from the Pomak cohort sequenced at a mean depth of 18.6x, using an identical sequencing, variant calling and quality control protocol. The proteomic phenotypes were transformed identically to MANOLIS. Single-point association analysis was performed using GEMMA and an identically calculated GRM. For burden replication, we specifically analysed the genes associated in MANOLIS in all conditions, and defined replication if a significant signal was detected in any of them. The COMT signal driven by rs4680 in MANOLIS (β=−0.373, σ=0.0405, *P*=3.5×10^−20^) could not be replicated due to COMT failing QC in Pomak. In five cases, associated MANOLIS variants and those tagged (r^2^>0.8) by them were monomorphic in the Pomak cohort (rs183455943, rs4778724, rs1053361963, rs200251994, rs186044494). 101/116 (87%) loci had at least one replicating variant.

### Definition of novelty

To assess whether a protein had been previously studied, we examined protein lists and summary statistics from five large published proteomics GWAS^15,30–33^. To determine novelty of genetic *cis* and *trans* association with proteins in our study, we first determined previously reported variants within a 2Mb window around the association peaks. Among these five GWAS, one^30^ performed stepwise conditional analysis to identify independent variants at associated loci, and three^15,31,33^ did LD-based detection of independent signals. We were unable to perform independent variant detection for the remaining study^32^ since no summary statistics were publicly available. We used GEMMA^24^ to perform association analysis using previously-reported independent variants as covariates. The association signals were declared novel if either there were no known signals in the 2Mb window, or the associations were still study-wide significant (*P*-value threshold: 7.45×10^−11^) after conditioning. For *trans* associations, we further annotated signals depending on whether they fell within highly pleiotropic genes that were associated with more than 1 protein in the current study and had evidence of additional associations in the literature (*KLKB1*, *ABO*, *APOE*, *FUT2*, *F12*), or whether they were independent of any *cis* signals in the vicinity. After this procedure, 58 *cis*-associated variants in 44 loci were either not within 1Mb or independent of a signal reported in previous proteomics GWAS. 22 *trans-*associated variants were both novel and independent from *cis* loci. 11 of these were not located within highly pleiotropic genes. For all loci annotated as provisionally novel using the above method, we queried the GWAS Catalog through the Ensembl REST API, as well as PhenoScanner^34^ in a 2Mb window around the lead SNP. Since proteomics GWAS signals are often designated generically in Ensembl, we additionally performed direct queries to the GWAS catalog REST API when phenotype descriptions were not specific enough. We manually investigated the list of signals in search of variants associated with the protein trait of interest. When such a variant was found, conditional analysis was performed and the novelty status updated accordingly. We further incorporated evidence from three association studies with signals that were not reported in the GWAS catalog ^35–37^. Using this method, there were thirty-nine proteins measured in this study for which we were not able to find evidence of previous studies of genetic associations. We find 8 *cis* loci harbouring 11 independent associated variants and one *trans* locus for 9 of these proteins.

### Variant consequences

Consequence was evaluated using Ensembl VEP^38^ for each variant with respect to any transcript of the *cis* gene for *cis-*associated variants and to the mapped gene for *trans-*associated variants. For *trans* associations, variants were manually mapped to any gene in a 1Mb window coding for known ligands or interactants when they were not contained within gene boundaries, as was the case for CXCL16 and LDLR. 16 replicating independent variants had a most severe consequence equal to or more severe than missense according to Ensembl VEP. For every variant, we extracted tagging SNVs at r^2^>0.8 using PLINK, however none of these tagging variants had a more severe consequence on the target gene than the independent variant. Similarly, we overlapped all independent variants with regulatory features using the Ensembl REST API. 35 variants in 29 loci overlapped with a regulatory feature. When extending the same analysis to variants with r^2^>0.8 with independent variants, 93 variants in 68 loci could be mapped to a regulatory feature.

### Colocalisation testing for eQTL overlap and pheWAS analysis

We perform colocalisation testing with eQTL data from the GTEX database^39^. First, for every signal, regions are extended 1Mb either side of every independent variant, and associations are conditioned on every other variant in the peak using GEMMA. For *cis* signals, expression information for the *cis* gene is extracted from the GTEX database over the same region. For *trans* signals, expression information is restricted to all genes located within a 2Mb region surrounding the variant. Then, for every variant/gene pair, we perform colocalisation testing using the fast.coloc function from the gtx R package (https://github.com/tobyjohnson/gtx). To account for multiple independent variants at the same locus, we perform conditional analysis on all independent variants except the one under consideration, and use these results as input for the colocalisation analysis. 60 (49%) independent variants in 45 (57%) *cis*-pQTLs co-localise with an expression quantitative trait locus (eQTL; identified in GTEx^40^) for the cis gene, in keeping with previous estimates for the proportion of pQTLs exerting their effects through transcriptional mechanisms^30^. In addition, we find that 29 (70%) *trans* signals co-localise with eQTLs for at least one gene in their vicinity (+/−1Mb).

We apply the same procedure to PhenoScanner data^34^, where eQTL data is replaced by the output of the PhenoScanner python utility 500kb either side of every independent variant. The colocalisation procedure is then repeated for every phenotype where at least one association is present in the region. Because such pheWAS results do not necessarily report weakly associated SNVs, the number of pheWAS variants to be colocalised can be small. We therefore modify the colocalisation script to handle the case where only one variant is present per phenotype. 82% of all signals co-localised with previous phenotypic associations (Supplementary Table 8). The proportion was similar for *trans* and *cis* loci, with 30 out of 37 co-localising with previous association signals in the latter case. As a proof of concept, we recapitulate strong evidence for colocalization of a progranulin-lowering trans-pQTL in the *SORT1-CELSR2* locus with increased risk of elevated cholesterol, angina pectoris, ischaemic heart disease and other cardio-damaging phenotypes. *trans*-pQTL for all the six proteins associated with an ABO region variant also colocalise with a range of blood and cardiovascular phenotypes, in line with a large body of evidence linking the ABO blood group with multiple diseases.

### Drug Target evaluation

For evaluating whether associated genes were drug targets, we used the OpenTargets^41^ and DrugBank^42^ databases. We accessed OpenTargets using the OpenTarget API. We converted the DrugBank XML file to flat files using the dbparser R package, and performed gene name matching using the USCS Gene Info database, downloaded May 6, 2019. 8 of the proteins for which a signal was detected at study-wide significance were targeted by drugs according to OpenTargets. This was true for 39 proteins when queried against the DrugBank database (Supplementary Table 9).

### Mouse models

We use the Ensembl REST API to extract mouse orthologs for all of the 109 genes whose protein for which genetic associations were found in our study. According to the IMPC API, KO experiments were reported for 26 of these orthologs, 18 of these having phenotypes associated with a p-value smaller than 1×10^−4^ (Supplementary Table 10).

### Two-Sample MR

We extracted variants characterized as independent signals by GCTA-COJO on a protein-by-protein basis across all *cis* and *trans* loci, and excluded novel variants without an rs-ID. For each remaining variant, we then considered summary statistics for all tagging positions with r^2^>0.8, merged the resulting data frame with the exposure dataset by rs-ID. All such records originating from all independent signals were then merged by protein and carried over to MR analysis using the MRBase R package. We excluded *trans* pleiotropic loci (*ABO*, *KLKB1*, *FUT2*, *APOE*, *F12*). MR was performed on a set of 127 cardiometabolic traits available in MRBase (Supplementary Table 3). Since all of our instruments involved a small number of variants (≤10), we used the inverse-variant weighted method, except for single-instrument analyses where we use the Wald ratio test, which consists of dividing the instrument-outcome by the instrument-exposure regression coefficient. We note that all of the GWAS summary statistics used in this analysis were not derived from WGS-based studies, and therefore several of our instruments were not found in these datasets and could not be used. An important caveat of our overlap-maximising approach is that we did not require overlapping variants to be lead variants in the outcome trait GWAS. This could potentially lead to false-positives for single-instrument tests if the variant is located at the shoulders of an association peak in the outcome trait GWAS. The future availability of population-scale association studies with WGS or WES will greatly enhance variant overlap compared to GWAS, and hence increase the power of MR analyses in proteomics.

We also leveraged summary statistics manually downloaded from recent large association studies for Chronic Kidney Disease^43^, blood lipids^44^, Atrial Fibrillation, Type-II Diabetes, Coronary Artery Disease, estimated glomerular filtration rate^45^, albuminuria^46^, and anthropometric traits^47^. Two proteins, PDL2 and TNFRSF10C, had both *cis* and *trans* associations where the *trans* did not fall in a highly pleiotropic gene. We performed 2-sample MR excluding *trans,* and found that no *cis* variant in TNFRSF10C could be found in selected external studies. This was due to the lead variant, 8:23108277 T/TG, being novel, and secondary variants, such as rs779159813, being very rare (MAF= 0.573%). For PDL2, the known *cis* signal driven by rs62556120 (β=0.416, σ=0.0276, *P*=2.92×10^−51^) was causally associated with increased risk of ulcerative colitis and inflammatory bowel disease (Supplementary Table 11), whereas in the *cis+trans* analysis, the addition of rs10935473 attenuates that signal, but drives a causal association with height.

### Significance thresholds

We calculate the significance thresholds by computing the effective number of variants, traits and analyses for every analysis requiring multiple-testing correction.

#### Single-variant analyses

The effective number of proteins was computed using the ratio of the eigenvalue variance to its maximum^48,49^:

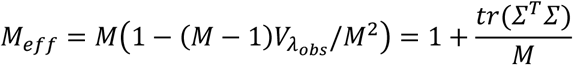

where 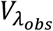 is the variance of the eigenvalues of the correlation matrix. For the *M* = 257 Olink phenotypes in this study, *M*_*eff*_ = 131.5, which we round to 132. The resulting p-value threshold is 7.45×10^−11^.

#### Rare-variant analyses

We report both single-point and rare variant burden signals, therefore increasing the multiple testing burden. The exact magnitude of this phenomenon being unknown, we performed a simulation study to compute the effective number of tests in case single-variant, variant-aggregation or both are reported in an association study. We find that reporting rare variant signals in combination with single-point signals at 5% and 1% MAF thresholds increased the multiple testing burden only marginally and by less than one order of magnitude (Supplementary Figure 7).

#### Two-sample MR

To correct for multiple testing in our MR analysis, we adjust p-values using FDR correction and examine significant results at an FDR of 0.05.

### Polygenic prediction

To examine predictive accuracy, we compute polygenic scores using PRSice 2^50^ with the Pomak as a target dataset. We assess scores between 1×10^−4^ and study-wide significance with intervals of 1×10^−10^. We further apply three MAF thresholds, at 0.05, 0.01 and MAC=10, which produces three best scores per protein. We find allele frequency thresholds not to have an appreciable influence on predictive power. The scores that achieve high accuracy (r^2^>0.05) in predicting Pomak protein levels all involve stringent thresholds (*P*<1×10^−6^, with the majority at *P*<1×10^−9^, Supplementary Table 12). To evaluate the predictive power of proteins for complex disease, we compute all scores between 1×10^−6^ and study-wide significance with the same interval. We then run a logistic regression of 86 UK Biobank ICD codes and self-reported traits (Supplementary Table 13) on all such scores on a protein-by-protein basis for 47 proteins that achieved r^2^>0.5 in the previous step. Sex, age, Qualification, Smoking status and BMI as well as 10 principal components are also added to the model. In order to accurately assess effect sizes, we select the most predictive score for each protein where at least one score threshold meets the Bonferroni-corrected P-value threshold (47 proteins, 64 effective phenotypes, *P*<1.66×10^−5^) in the Wald test for variable contribution. We then run the same model as before, with only this score and covariates as predictors. We removed associations between 3 proteins (CTRC, ICAM2, SELE) and deep vein thrombosis and/or pulmonary embolism due to the association being entirely driven by *ABO* variants.

In order to validate our approach for variable selection, we pooled all scores for all proteins in a single model of self-reported high cholesterol, and used elastic net regression to shrink coefficients for non-informative predictors. We run 10-fold cross-validation with a 20% hold-out sample to optimize the lambda value, and run 11 models using different values of alpha, from 0 (ridge regression) to 1 (lasso) at intervals of 0.1. We use a value of lambda one standard deviation away from the minimal value to increase shrinkage. Overall, the performance was comparable across models, but small values of alpha yielded a better predictive performance on the holdout set. An alpha value of 0.1 yielded almost no loss of AUC compared to ridge regression, but shrunk almost all coefficients to 0 (Supplementary Figure 8). Four protein scores (CHI3L1, PECAM1, SELE and GRN) were shrunk to a value greater than 0, confirming results of the manual variable selection procedure.

## Acknowledgements

We thank the residents of the Pomak and Mylopotamos villages for taking part. The MANOLIS study is dedicated to the memory of Manolis Giannakakis, 1978–2010. This work was funded by the Wellcome Trust [098051] and the European Research Council [ERC-2011-StG 280559-SEPI]. The GATK3 program was made available through the generosity of the Medical and Population Genetics program at the Broad Institute, Inc. We thank the Human Genetics DNA Pipelines and Human Genetics Informatics departments at the Wellcome Sanger Institute for performing sequencing and variant calling. This study has been conducted using the UK Biobank Resource (project ID 10205).

## Author Contributions

Sample collection and phenotyping: ET, MK, GD, EZ

Phenotype transformation and quality control: AG, YCP, GP, LS, NWR

Association analysis: AG, YCP

Software development: AG, DS

Bioinformatics: AG, SN, IF, AB

Simulation analysis: TB

Study design and supervision: EZ

Manuscript writing: AG, IF, EZ

