## Supplementary Material for "Whole genome sequencing analysis of the cardiometabolic proteome"

**Supplementary Figures**

**Supplementary Figure 1: *cis*-association within the *CTSH* gene. Independent variants as defined by COJO[1] are highlighted by crosses.**

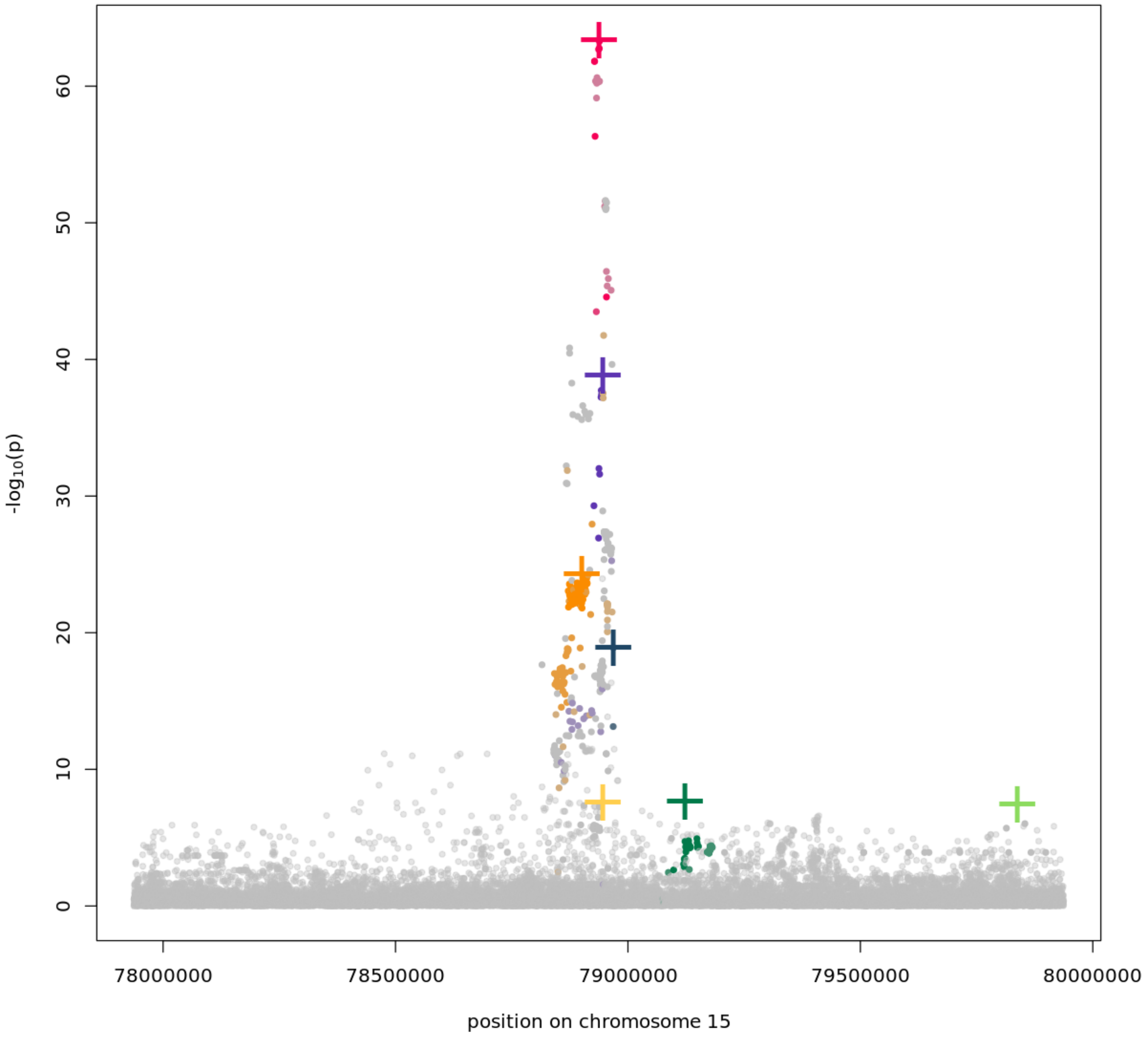

**Supplementary Figure 2: Effect size according to log-MAF for independent variants at pQTLs discovered in this study.** Question marks denote variants not present in the replication dataset and for which no LD-based proxy ($r^2$>0.8) could be found, crosses indicate variants that were tested but did not pass the replication significance threshold.

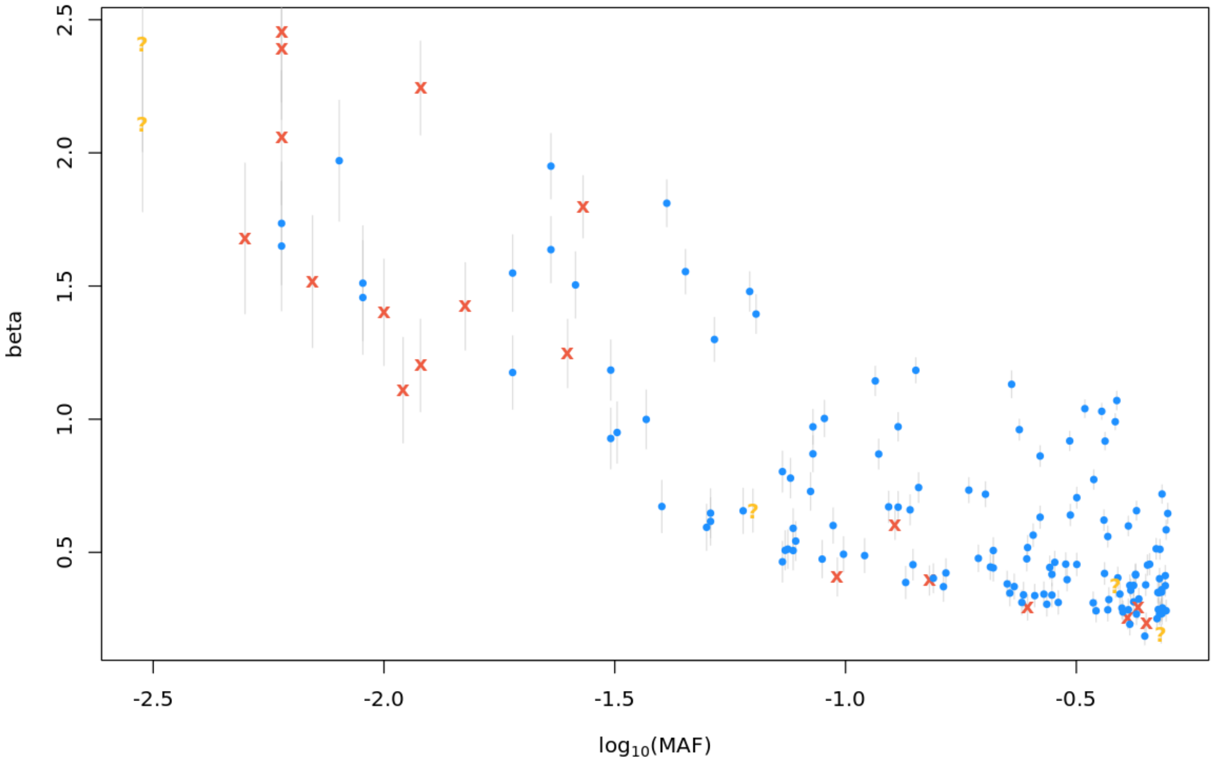

**Supplementary Figure 3: Variance explained in proteomic traits compared with 37 non-proteomic traits (Supplementary Table 15) in the same cohort using the same association protocol.**

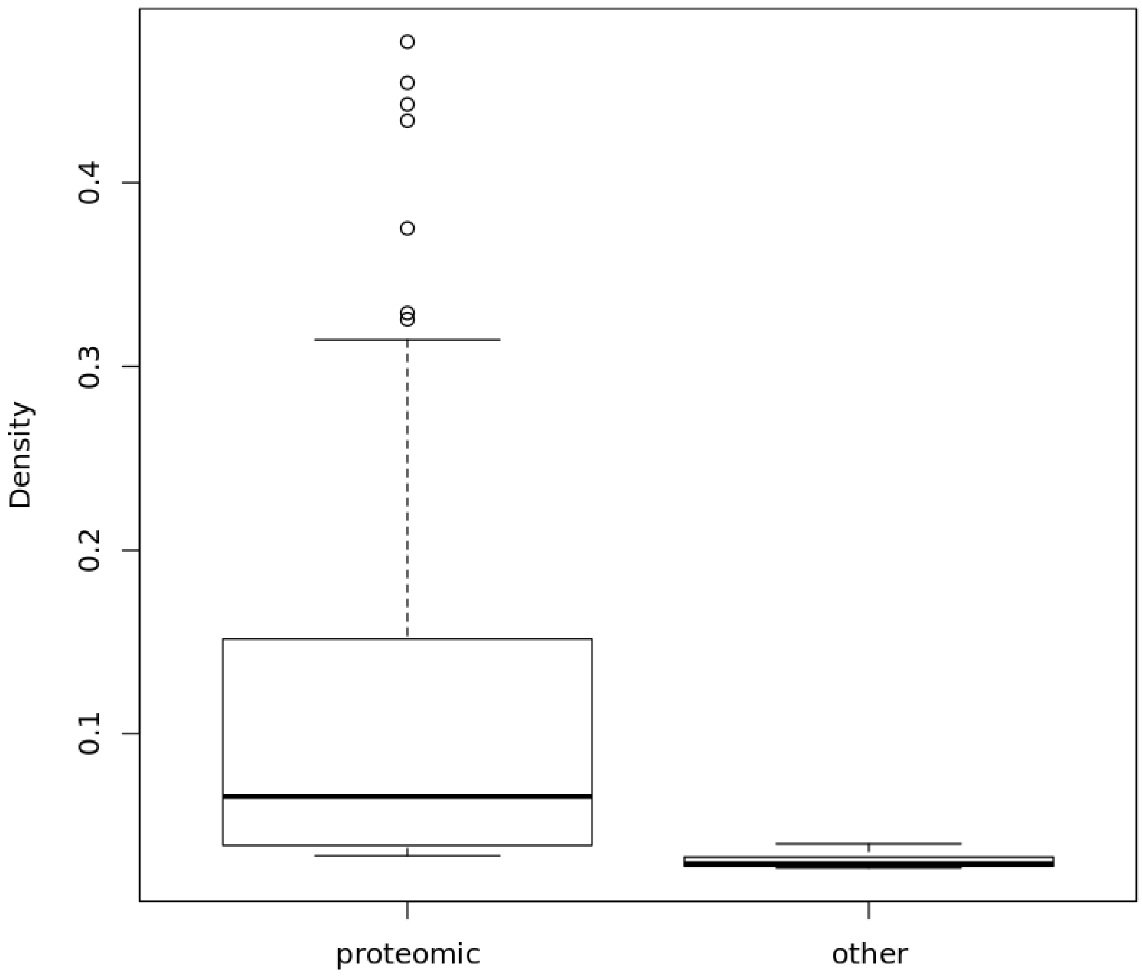

**Supplementary Figure 4: Distance to gene boundary as defined by Ensembl REST API for *cis* independent variants.**

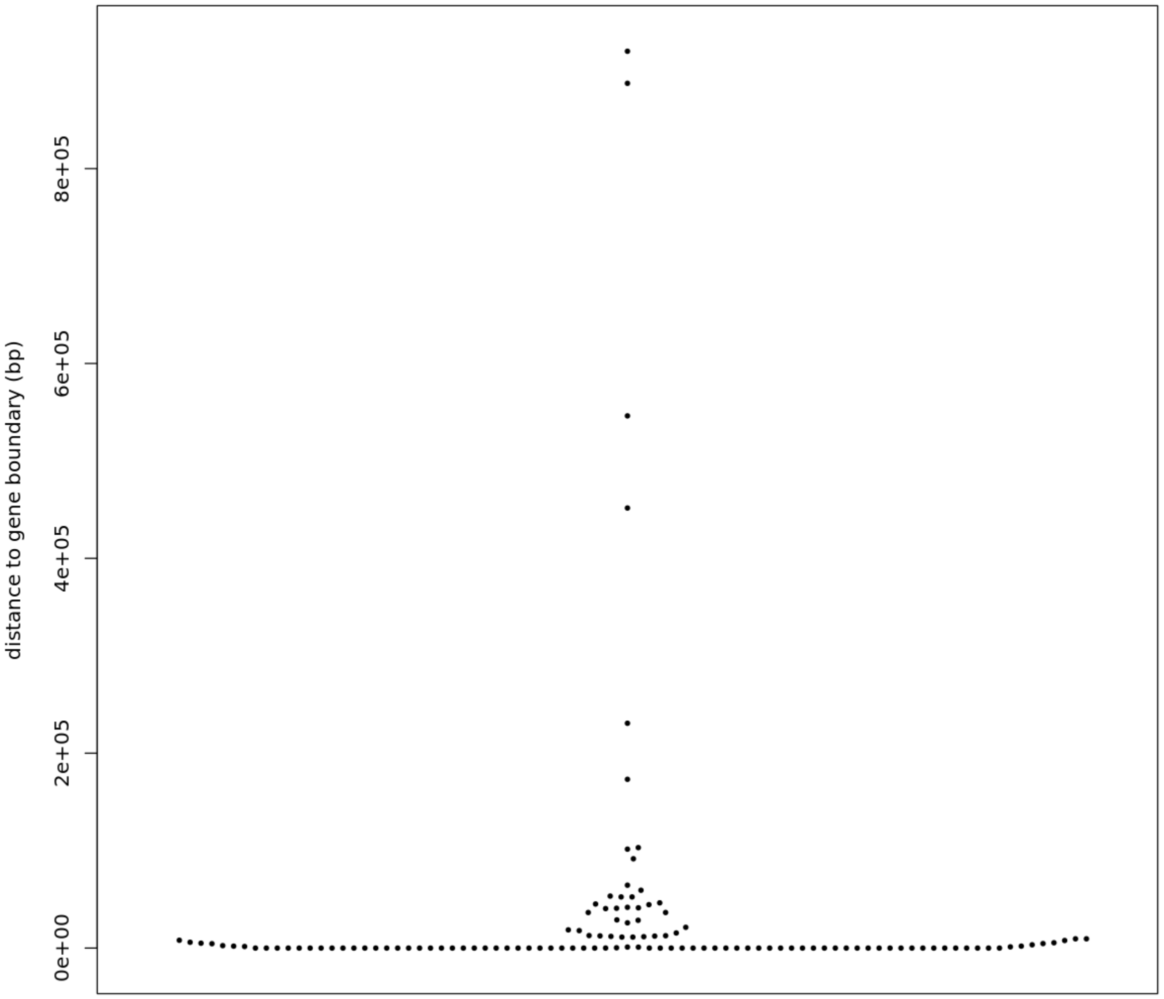

**Supplementary Figure 5: Significant burden associations across all analyses.** Numbers and colour scale represent significance on the -log10 scale. Columns are variant selection methods (see Methods), rows are the proteins for which these significant *cis*-acting associations were found.

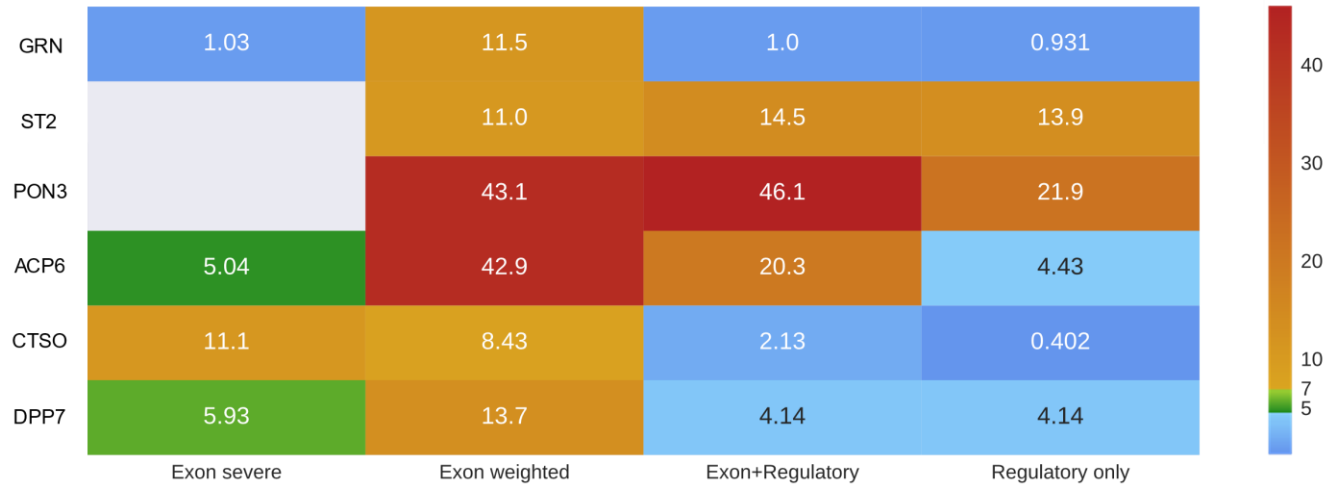

**Supplementary Figure 6: Evaluation of effective number of tests and significance thresholds.** Reporting single-point (a,d), sliding-window rare variant burden test (b, e) and both (c, f), using two different simulation models (a, b, c and d, e, f, respectively), on chromosome 11, across 1,000 simulations.

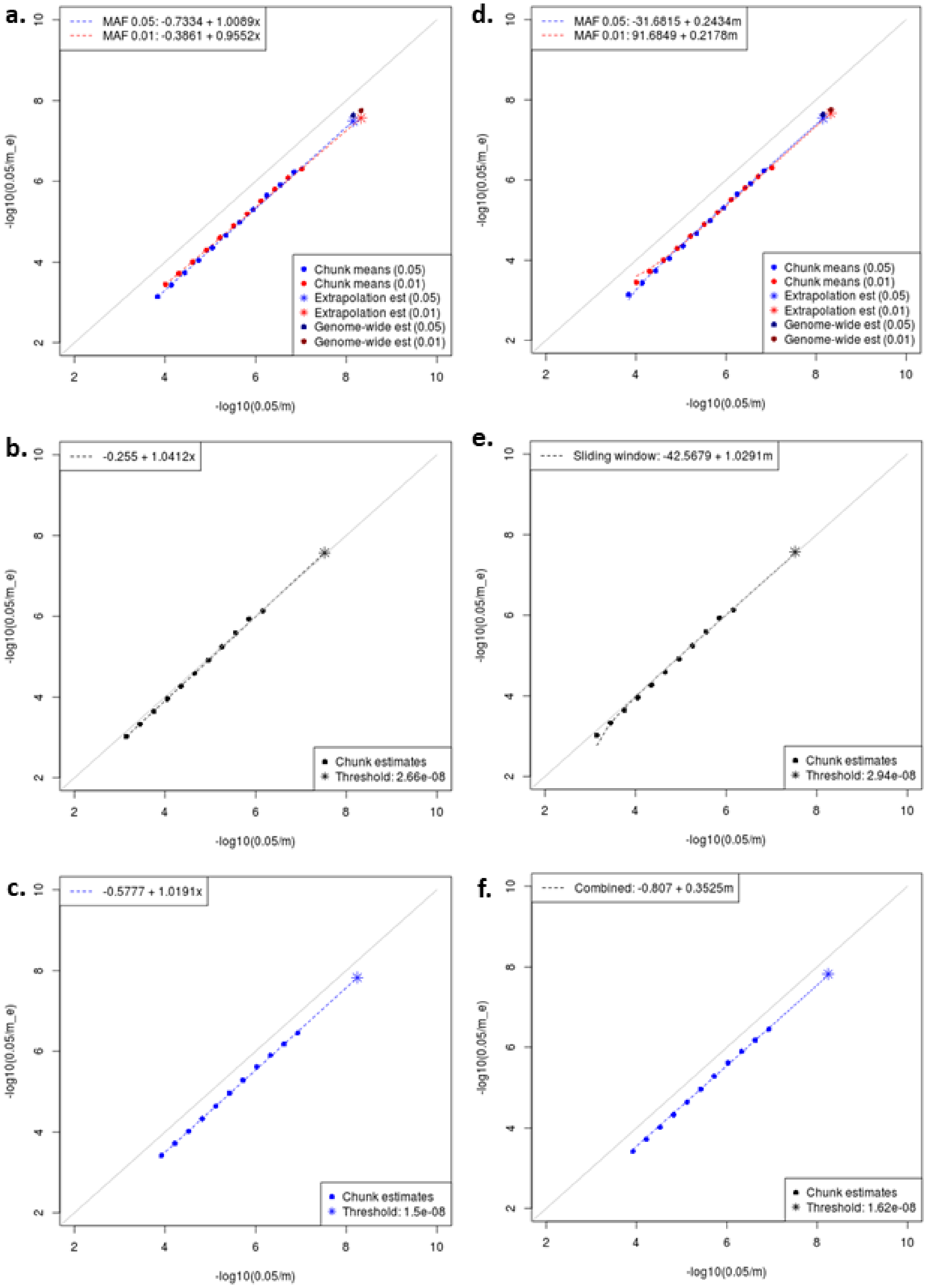

**Supplementary Figure 7. Elastic net model of high cholesterol.** Top left: Precision-recall curves for models with different values of alpha on the holdout set. The ridge model curve (alpha=0) is coloured by lambda value according to the scale on the right of the plot. All other models are drawn on a purple to yellow scale, with yellow corresponding to alpha=0.1 and purple to alpha=1. Top right: Area under the curve for the optimal lambdas + 1 standard error, according to alpha, for predictions on the holdout set. Bottom left: Mean-square error estimate and standard error according to lambda, for alpha=0.1, on the training set. The two vertical lines indicate the minimal lambda and the minimal lambda + 1 standard error, respectively. Bottom left: Coefficient trajectory as a function of log(lambda) for the training set, with the top 4 coefficients in absolute value highlighted. The model coefficients fit on the entire dataset are given in the supplementary Text.

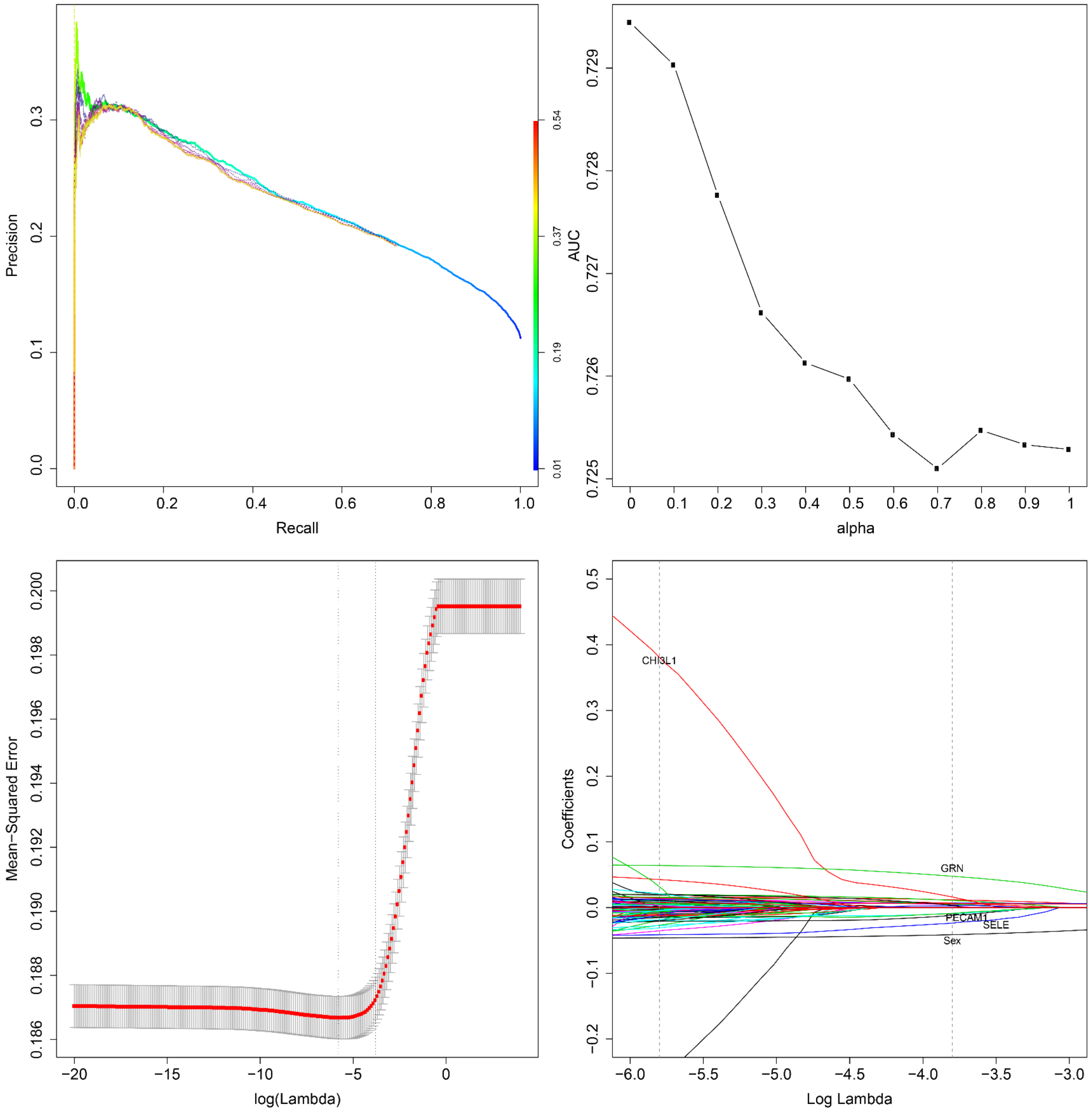

**Supplementary Text**

### *Additional notable pQTL signals*

The *PLAUR* missense variant rs4760 is associated with decreased levels of TNFRSF10C (TNF receptor superfamily member 10c) (MAF=13%, β=-0.96, σ=0.054, *P*=7.31x$10^{-52}$). *PLAUR* codes for the urokinase receptor (uPAR). Urokinase is an essential thrombolytic agent and acts as an invasion-promoting protein in several types of cancer[2] through blocking efferocytosis and phagocytosis, notably in apoptotic cardiocytes[3]. This variant has previously been associated with decreased levels of the TRAIL apoptosis-inducing ligand[4], TNFSF10C. TRAIL has been shown to induce overexpression of urokinase, and uPAR acts as a "don't eat me" signal for apoptotic cells. This *trans*-pQTL finding could indicate that impairment of the urokinase receptor is linked to an oversensitivity to TRAIL signalling, leading to decreased levels of both TRAIL and its receptor.

rs10886430, an intronic *GRK5* variant, is associated with decreased CCL17 levels (MAF=9.9%, β=-0.493, σ=0.0656, *P*=3.72x$10^{-13}$). CCL17 restrains regulatory T cell homeostasis to promote atherosclerosis through binding to CCR4 and other receptors[5]. Acting through G protein-coupled chemokine receptors, ACKR1 ligands such as CCL17 can induce activation and migration of leucocyte subsets into the vessel wall, and play a pathogenic role during atherosclerosis development[6]. *GRK5* codes for a G protein-coupled receptor kinase, which desensitises activated G protein-coupled receptors through phosphorylation and subsequent binding of arrestin. This *trans*-pQTL finding could indicate a phosphorylation activity of GRK5 on one of the G-coupled receptors of CCL17 such as CCR4 or CCR8. rs10886430 is not a significant eQTL for any gene in any tissue[7].

rs144846334, an intergenic variant upstream of *SLC10A2*, is associated with decreased levels of EPCAM (β=-0.779, σ=0.0738, MAF=7.6%, *P*=3.52x$10^{-23}$, *trans*-pQTL). rs144846334 is in strong LD ($r^2$>0.8) with the *SLC10A2* missense variant rs56398830 (β=-0.77, σ=0.0730, *P*=5.42x$10^{-23}$). *SLC10A2* plays a key role in sodium-dependent intestinal bile salt reuptake, and variants in this gene have been associated with numerous phenotypes such as gallbladder diseases, venous thromboembolism, LDL and HDL cholesterol. *SLC10A2* knockout mice have decreased intestinal cholesterol absorption, abnormal bile salt levels and steatorrhea. Loss of *EPCAM* function, is causal for congenital tufting enteropathy[8], a severe sodium-losing diarrheal disorder presenting in the neonatal period.

rs1309620228, a start lost variant (MAF=0.2%, β=-2.21, σ=0.410, *P*=1.21x$10^{-7}$), and rs556026695, a splice donor variant (MAF=0.2%, β=-2.36, σ=0.457, *P*=3.63x$10^{-7}$) drive a *cis*-RV-pQTL for CTSO (*P*=7.94x$10^{-12}$). Both variants are present at much lower frequencies in cosmopolitan populations (MAC=1 in TOPMed for rs1309620228; MAC=1 in gnomAD and TOPMed for rs556026695). A third variant, the splice region variant rs763411023, is included but its contribution to the burden is small (*P*=0.066). The *CTSO* gene codes for cathepsin O, a cysteine protease with unclear function. Cathepsin O is ubiquitously expressed, and is involved in normal cellular protein degradation and turnover[9]. In mice, mutations in the *Ctso* gene have been associated with decreased bilirubin and aspartate transaminase levels. rs11722604, an intronic *CTSO* variant, has previously been associated with increased adiponectin in an East Asian cohort. This rare variant burden signal replicates in Pomak (*P*=4.01x$10^{-24}$ in the exon weighted analysis), and is entirely driven by the missense variant rs1013059201.

### *Joint model of high cholesterol using CHI3L1 and PECAM1*

To quantify cumulative contributions of predictive proteins to hypercholesterolemia risk, we perform a joint logistic model of high cholesterol using CHI3L1 and PECAM1 scores, including clinical and genetic covariates as predictors. A third protein, GRN, was also significantly associated with hypercholesterolemic outcomes, however we did not include it in our joint model due to it being driven by a well-known association at the *SORT1* locus. The full model is reported below:

```
Call:
glm(formula = "high_cholesterol~.", family = binomial(link = "logit"), 
    data = m)

Deviance Residuals: 
    Min       1Q   Median       3Q      Max  
-1.6071  -0.5381  -0.3845  -0.2570   3.1113  

Coefficients:
                                   Estimate Std. Error  z value Pr(>|z|)    
(Intercept)                      -9.1603766  0.0573380 -159.761  < 2e-16 ***
Sex2                             -0.5160839  0.0107381  -48.061  < 2e-16 ***
Smoking_status1                   0.1430716  0.0112802   12.683  < 2e-16 ***
Smoking_status2                   0.2415334  0.0185070   13.051  < 2e-16 ***
Age_when_attended_assessment_centre  0.0899566  0.0007776  115.689  < 2e-16 ***
Qualifications2                   0.0985557  0.0170953    5.765 8.16e-09 ***
Qualifications3                   0.1647312  0.0134691   12.230  < 2e-16 ***
Qualifications4                   0.2193386  0.0238572    9.194  < 2e-16 ***
Qualifications5                   0.1889407  0.0186205   10.147  < 2e-16 ***
Qualifications6                   0.1748300  0.0204647    8.543  < 2e-16 ***
PC1                               0.0011252  0.0001022   11.014  < 2e-16 ***
PC2                              -0.0034697  0.0001912  -18.151  < 2e-16 ***
PC3                               0.0056005  0.0003398   16.482  < 2e-16 ***
PC4                              -0.0021145  0.0004537   -4.661 3.15e-06 ***
PC5                               0.0002571  0.0007098    0.362   0.7172    
PC6                              -0.0005228  0.0011003   -0.475   0.6347    
PC7                              -0.0001937  0.0010362   -0.187   0.8517    
PC8                              -0.0020216  0.0011046   -1.830   0.0672 .  
PC9                               0.0006980  0.0011411    0.612   0.5407    
PC10                              0.0062680  0.0012046    5.203 1.96e-07 ***
BMI                               0.0705270  0.0010475   67.327  < 2e-16 ***
PECAM1                           -0.7148646  0.0783299   -9.126  < 2e-16 ***
CHI3L1                            0.7255380  0.1181090    6.143 8.10e-10 ***
---
Signif. codes:  0 '***' 0.001 '**' 0.01 '*' 0.05 '.' 0.1 ' ' 1

(Dispersion parameter for binomial family taken to be 1)

    Null deviance: 278033  on 395511  degrees of freedom
Residual deviance: 252115  on 395489  degrees of freedom
  (91897 observations deleted due to missingness)
AIC: 252161

Number of Fisher Scoring iterations: 5
```

Both protein scores contribute to the model. The nested model excluding protein scores is given below:

```
Call:
glm(formula = "high_cholesterol~.", family = binomial(link = "logit"), 
    data = m[, -c("PECAM1", "CHI3L1"), with = F])

Deviance Residuals: 
    Min       1Q   Median       3Q      Max  
-1.6277  -0.5384  -0.3851  -0.2575   3.1026  

Coefficients:
                                   Estimate Std. Error  z value Pr(>|z|)    
(Intercept)                      -9.1479772  0.0570438 -160.368  < 2e-16 ***
Sex2                             -0.5159580  0.0107361  -48.058  < 2e-16 ***
Smoking_status1                   0.1428811  0.0112779   12.669  < 2e-16 ***
Smoking_status2                   0.2407488  0.0185043   13.010  < 2e-16 ***
Age_when_attended_assessment_centre  0.0899048  0.0007774  115.654  < 2e-16 ***
Qualifications2                   0.0990247  0.0170914    5.794 6.88e-09 ***
```

```
Qualifications3                          0.1648836  0.0134662   12.244  < 2e-16 ***
Qualifications4                          0.2199415  0.0238540    9.220  < 2e-16 ***
Qualifications5                          0.1889738  0.0186179   10.150  < 2e-16 ***
Qualifications6                          0.1753056  0.0204601    8.568  < 2e-16 ***
PC1                                      0.0011130  0.0001020   10.910  < 2e-16 ***
PC2                                     -0.0034758  0.0001911  -18.189  < 2e-16 ***
PC3                                      0.0055479  0.0003397   16.333  < 2e-16 ***
PC4                                     -0.0023560  0.0004529   -5.203 1.97e-07 ***
PC5                                     -0.0001256  0.0007084   -0.177   0.8592    
PC6                                     -0.0005892  0.0010996   -0.536   0.5921    
PC7                                     -0.0001620  0.0010354   -0.156   0.8757    
PC8                                     -0.0020606  0.0011037   -1.867   0.0619 .  
PC9                                      0.0004355  0.0011402    0.382   0.7025    
PC10                                     0.0062708  0.0012040    5.209 1.90e-07 ***
BMI                                      0.0704985  0.0010472   67.320  < 2e-16 ***
---
Signif. codes:  0 '***' 0.001 '**' 0.01 '*' 0.05 '.' 0.1 ' ' 1

(Dispersion parameter for binomial family taken to be 1)

    Null deviance: 278033  on 395511  degrees of freedom
Residual deviance: 252235  on 395491  degrees of freedom
  (91897 observations deleted due to missingness)
AIC: 252277

Number of Fisher Scoring iterations: 5
```

The likelihood ratio test was performed using the lrtest function in the lmtest package, and produces

### *Coefficients of the elastic net model*

The coefficients for the elastic net model described in the text, fit on the entire dataset for more accurate parameter estimation, are given in the table below. The qualifications variables are a dummy coding of the UK Biobank field 6138 using data coding 100305 where variable levels have been turned into labels, and the smoking status variables are a dummy coding of the UK Biobank field 20116 using data coding 90. For protein genetic risk scores, the P-value threshold is included in parentheses.

| Variable group | Variable | Effect |
|---|---:|---:|
| | (Intercept) | 0.587814192600955 |
| | Sex | -0.0413162080627943 |
| Smoking status (dummy variables, vs. never smoked) | Previous Smoker | 0.0118989443411119 |
| | Current Smoker | 0.0111962448743365 |
| | Age | 0.00684558775742107 |
| Qualifications (dummy vaiables, vs. University degree) | O levels/GCSEs or equivalent | 0.00394959865872145 |
| | NVQ or HND or HNC or equivalent | 0.00895163532749535 |
| | Other professional qualifications eg: nursing, teaching | 0.00528342948315935 |
| Principal components (PCs) | PC1 | 7.47342507105865e-05 |
| | PC2 | -0.000213788142332847 |
| | PC3 | 0.00044419244443404 |
| | PC10 | 3.83494557705365e-05 |
| | BMI | 0.00606310791867212 |
| Protein scores | CHI3L1 ($P<9.25\text{x}10^{-7}$) | 0.0199821126019553 |
| | GRN ($P<7.45\text{x}10^{-11}$) | 0.0453636405134527 |
| | GRN ($P<4.57\text{x}10^{-9}$) | 0.00589023195891766 |
| | PECAM1 ($P<1.95\text{x}10^{-7}$) | -0.01608456535475 |
| | SELE ($P<1.37\text{x}10^{-9}$) | -0.00651275802010374 |
| | SELE ($P<8.16\text{x}10^{-8}$) | -0.00011048595528366 |
| | SELE ($P<8.87\text{x}10^{-8}$) | -0.0174515150666494 |
| | SELE ($P<1.77\text{x}10^{-7}$) | -0.00352572640455521 |
